# Morphognostic honey bees communicating nectar location through dance movements

**DOI:** 10.1101/2020.03.14.992263

**Authors:** Thomas E. Portegys

**Affiliations:** Kishwaukee College, Malta, Illinois USA

**Keywords:** Honey bee foraging dance, Morphognosis, artificial animal intelligence, artificial neural network, machine learning, artificial life, cellular automaton

## Abstract

Honey bees are social insects that forage for flower nectar cooperatively. When an individual forager discovers a flower patch rich in nectar, it returns to the hive and performs a “waggle dance” in the vicinity of other bees that consists of movements communicating the direction and distance to the nectar source. The dance recruits “witnessing” bees to fly to the location of the nectar to retrieve it, thus cooperatively exploiting the environment. Replicating such complex animal behavior is a step forward on the path to artificial intelligence. This project simulates the bee foraging behavior in a cellular automaton using the Morphognosis machine learning model. The model features hierarchical spatial and temporal contexts that output motor responses from sensory inputs. Given a set of bee foraging and dancing exemplars, and exposing only the external input-output of these behaviors to the Morphognosis learning algorithm, a hive of artificial bees can be generated that forage as their biological counterparts do. A comparison of Morphognosis foraging performance with that of an artificial recurrent neural network is also presented.

## 1. Introduction

Honey bees, *apis mellifera*, are fascinating social insects. They are also smart, even able to count and add (Fox 2019). However, it is their ability to communicate symbolically in the form of a “waggle dance” indicating the direction and distance to a nectar source that is truly astonishing (Chittka and Wilson 2018; Nosowitz 2016; Schürch et al. 2019) especially considering that the use of symbols is rare even in more neurologically complex animals. In 1973, the Austrian scientist Karl von Frisch was awarded the Nobel Prize for his research on this behavior (1967).

The waggle dance is a figure-eight movement, done by a bee in the presence of other bees in the hive after discovering a food source at a locale outside the hive, which recruits bees to forage at the indicated location, thus acquiring more food than solitary foraging would otherwise. The dance includes information about the direction and distance to the goal. Distance is indicated by the length of time it takes to make one figure-eight circuit. Direction of the food source is indicated by the direction the dancer faces during the straight portion of the dance when the bee is waggling. This indicates the angle from the sun to the goal. For example, if the bee waggles while facing straight upward, then the food source may be found in the direction of the sun.

This paper describes artificial honey bees that gather nectar and perform an analog of the waggle dance. It employs a general machine learning system, *Morphognosis*, which acquires behaviors by example and enables an artificial organism to express those behaviors. It will be shown that nectar foraging is a daunting task for unaided machine learning methods, but with the support of the spatial and temporal contextual information provided by Morphognosis, it can be accomplished.

As a disclaimer, it should be noted that this project is not intended to offer new or additional findings about honey bees. Neither does it simulate many honey bee behaviors. For example, honey bees use the sun for navigation. This is not simulated.

Honey bees have been the focus and inspiration for a number of simulation initiatives:

- Food source recruitment in honey bees (Dornhaus et al. 2006).
- Detailed colony behavior, including nectar foraging (Betti et al. 2017).
- Swarming and group behavior algorithms (Karaboga and Akay 2009).
- Flight neural network (Cope et al. 2013).
- Visual system neural network (Roper et al. 2017).
- Odor learning circuits (MaBouDi et al. 2017).
- Spiking neural network that reacts to nectar (Fernando and Kumarasinghe 2015).

The food source recruitment simulation uses a state machine, following rules developed from experiments with bees, to control bee foraging behavior. The colony simulation allows a user to observe how bees are affected by various environmental conditions, such as weather. Algorithms for a number of group behaviors, optimal foraging strategies among them, are cited in the Karaboga and Akay paper. The other projects simulate bee-specific neural mechanisms. For example, the odor learning project found that simulated honey bees lacking mushroom bodies, the insect equivalent of the cerebral cortex, may still be able to learn odors. The spiking neural network measures how an abstracted model of a bee’s nervous system reacts to nectar-related stimuli.

The above simulations are designed for specific bee functionalities, and accordingly are inapplicable outside of their domains of operation. In contrast, the intended contribution of this project is to replicate honey bee behavior with a general purpose model that learns from external observations and which is applicable to many behavioral tasks, not just the honey bee foraging task.

A number of years ago I explained to a coworker how my dissertation program (Portegys 1986), a model of instrumental/operant conditioning, could learn various tasks through reinforcement. He then asked me how “smart” it was. I put him off, not having a ready answer. He persisted. So I blurted out that it was as smart as a cockroach (which it is not). To which he replied, “Don’t we have enough *real* cockroaches?” Fast forward to this project. Don’t we have enough *real* honey bees? (Although, come to think of it, maybe we don’t (Oldroyd 2007)!)

The point of this story is that the question of why anyone should work on artificial animal intelligence is, at least on the surface, a reasonable one, given our species unique intellectual accomplishments. Thus, historically, AI has mostly focused on human-like intelligence, for which there are now innumerable success stories: games, self-driving cars, stock market forecasting, medical diagnostics, language translation, image recognition, etc. Yet the elusive goal of artificial general intelligence (AGI) seems as far off as ever. This is because these success stories lack the “general” property of AGI, operating as they do within narrow, albeit deep, domains. A language translation application, for example, does just that and nothing else.

Anthony Zador (2019) expresses this succinctly: “We cannot build a machine capable of building a nest, or stalking prey, or loading a dishwasher. In many ways, AI is far from achieving the intelligence of a dog or a mouse, or even of a spider, and it does not appear that merely scaling up current approaches will achieve these goals.”

I am in the camp that believes that achieving general animal intelligence is a necessary, if not sufficient, path to AGI. While imbuing machines with abstract thought is a worthy goal, in humans there is a massive amount of ancient neurology that underlies this talent.

Hans Moravec put it thusly (1988): “Encoded in the large, highly evolved sensory and motor portions of the human brain is a billion years of experience about the nature of the world and how to survive in it. The deliberate process we call reasoning is, I believe, the thinnest veneer of human thought, effective only because it is supported by this much older and much more powerful, though usually unconscious, sensorimotor knowledge. We are all prodigious Olympians in perceptual and motor areas, so good that we make the difficult look easy. Abstract thought, though, is a new trick, perhaps less than 100 thousand years old. We have not yet mastered it. It is not all that intrinsically difficult; it just seems so when we do it.”

So how should we proceed? Emulating organisms at the level of neurons (whole-brain emulation) is a possible approach to understanding animal intelligence. However, efforts to do this with the human brain have met with little success (Yong 2019). Scaling down to mice is an option. The human brain dwarfs the mouse brain, but even mouse brains are daunting: a cubic milliliter of mouse cortex contains 900,000 neurons and 700,000,000 synapses (Braitenberg and Schüz 1998). At much a simpler scale, years have been spent studying the relationship between the connectome of the nematode C. elegans (Wood 1988), with only 302 neurons, and its behaviors, but even this creature continues to surprise and elude full understanding. Despite this, some researchers believe that it is now feasible for the whole-brain approach to be applied to insects such as the fruit fly, with its 135,000 neurons (Collins 2019). Partial brain analysis is also an option. For example, the navigation skills of honey bees are of value to drone technology. Fortunately, it appears that the modular nature of the honey bee brain can be leveraged to replicate this skill (Nott 2018).

Another issue with emulation is the difficulty of mapping the relationship between neural structures and behaviors (Krakauer et al. 2017; Yong 2017). For AI, this is a key aspect, as behavior is the goal. Nature is a blind tinkerer, unbound by design rules that appeal to humans. For example, despite the enthusiasm following the mapping of the human genome, the mechanisms by which genes express proteins, and thus phenotypes, is not as modular as hoped for. Rather, it is extraordinarily complex (Boyle et al. 2017; Wade 2001). When blindly copying a natural system into an artificial one, artifacts and quirks left over by evolution can introduce unnecessary complexity.

The field of artificial life (Alife) offers another path to AGI. This path starts with simulating life, and letting evolution optimize artificial organisms to achieve intelligence as a fitness criteria. For example, Schöneburg’s (2019) “alternative path to AGI”, sees intelligence emerging from *holobionts*, which form cooperating collectives of artificial agents.

Morphognosis carries on the trend set by artificial neural networks to abstractly model neurological computing functions. However, the approach is primarily to simulate at the behavioral level. Considering the vastly different “clay” that biological and computing systems are built with, cells vs. transistors and software, behavioral simulation seems a good place to converge. The famous Turing Test (1950) follows this line of thought.

Morphognosis comprises an artificial neural network (ANN) enhanced with a framework for organizing sensory events into hierarchical spatial and temporal contexts. Nature has hard-wired knowledge of space and time into the brain as way for it to effectively interact with the environment (Bellmund et al. 2018; Buffalo 2015; Hainmüller and Bartos 2018; Lieff 2015; Vorhees and Williams 2014). These capabilities are modeled by Morphognosis. Interestingly, in humans spatial mapping cells called grid cells appear to be capable of representing not only spatial relationships, but non-spatial multidimensional ones, such as the relationships between members of a group of people (Bruner et al. 2018; Tavares et al. 2015).

The bee dancing behavior, as a sequential process, has temporal components. For example a bee must remember a past event, the existence of surplus nectar in a flower, and use that information to perform a dance that indicates both direction and distance to the nectar. In addition, bees that observe a dance must internally persist the distance signal and use it to measure how far to fly.

Sequential processes are type of task that recurrent artificial neural networks (RNNs) have been successfully applied to (Elman 1990; Hochreiter and Schmidhuber 1997). However, RNNs do not support spatial information. RNNs maintain internal feedback that allow them to retain state information within the network over time. This contrasts with Morphognosis, where the input itself contains temporal state information.

Morphognosis was partly inspired by some what-if speculation. In simpler animals, the “old” brain (amygdala, hypothalamus, hippocampus, etc.) deals more directly with a less filtered here-and-now version of the environment. Considering nature’s penchant for repurposing existing capabilities, might it be that in more complex animals a purpose of the neocortex, sitting atop the old brain and filtering incoming sensory information, is to track events from distant reaches of space and time and render them, as though near and present, to the old brain whose primal functions have changed little over time? In this project, an ANN plays the role of the old brain, and Morphognosis is the counterpart to the neocortex.

I have previously conducted research into a number of issues that differentiate conventional AI from natural intelligence. These include context, motivation, plasticity, modularity, instinct, and surprise (Portegys 2007, 2010, 2013, 2015). Morphognosis, in particular, has been previously applied to the task of nest-building by a species of pufferfish (Portegys 2019).

To date, including the honey bee project, Morphognosis has been implemented as a cellular automaton (Toffoli and Margolus 1987; Wolfram 2002), as the rules that it develops while learning are ideally captured in a grid structure. Conceptually, however, Morphognosis is not tied to the cellular automaton scheme.

The next section describes Morphognosis and details of the behavior and implementation of the honey bees. A section with the results of testing pertinent variables follows. Finally, a comparison of the performance of a recurrent neural network on the foraging task is presented (see LSTM performance).

## 2. Description

This section first briefly describes the Morphognosis model. The honey bee behavior and implementation are described next.

### 2.1. Morphognosis overview

*Morphognosis* (*morpho* = shape and *gnosis* = knowledge) aims to be a general method of capturing contextual information that can enhance the power of an artificial neural network (ANN). It provides a framework for organizing spatial and temporal sensory events and motor responses into a tractable format suitable for ANN training and usage.

Introduced with several prototype tasks (Portegys 2017), Morphognosis has also modeled the locomotion and foraging of the C. elegans nematode worm (Portegys 2018) and the nest-building behavior of a pufferfish (Portegys 2019). Morphognosis is a temporal extension of a spatial model of morphogenesis (Portegys et al. 2017).

#### 2.1.1. Morphognostics

The basic structure of Morphognosis is a cone of multi-dimensional sensory event vectors called a *morphognostic*, shown in Figure 1. At the apex of the cone are the most recent and nearby sensory events. Receding from the apex are less recent and possibly more distant events. A mobile animal can generate spatial movements which produce sensory inputs that are geographically encoded in the morphognostic, analogously to place cells in the hippocampus (Vorhees and Williams 2014).

**Figure 1 -.**
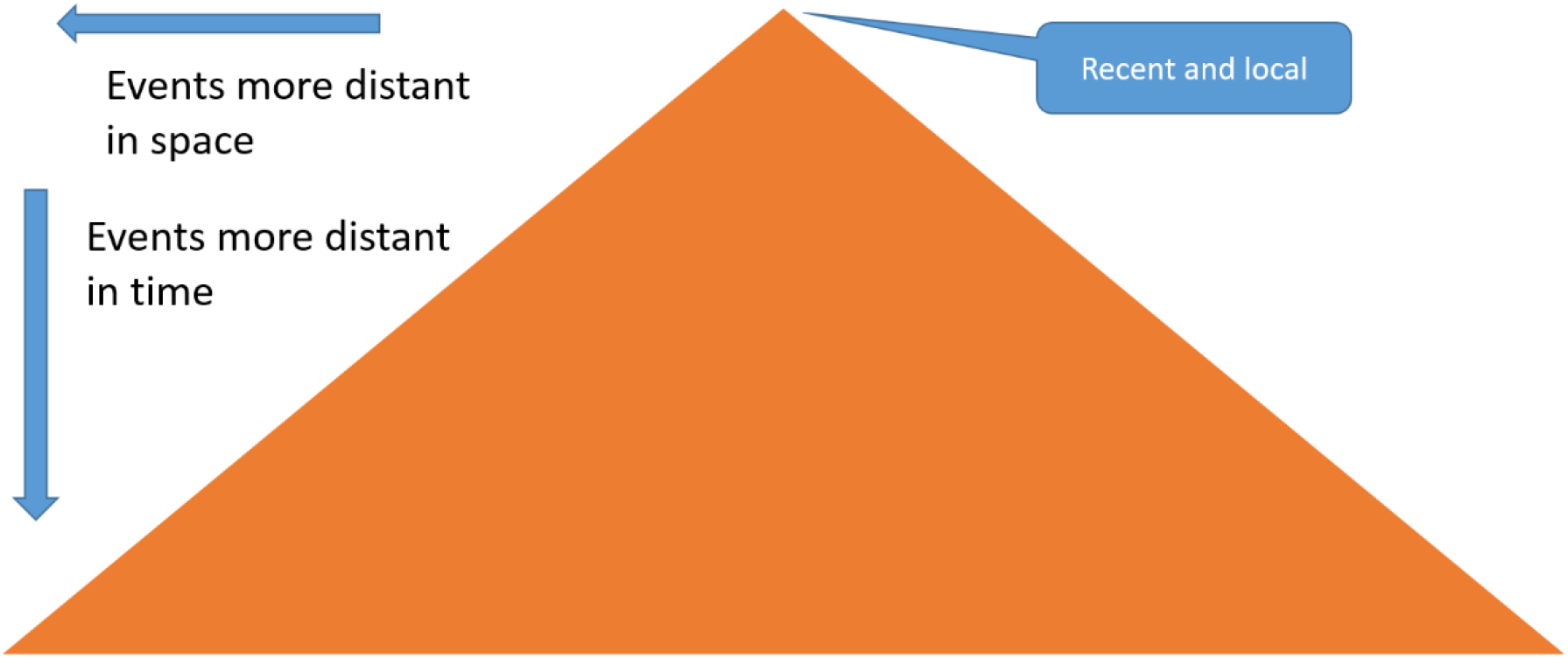
Morphognostic sensory event cone.

Sensory event vectors are aggregated in the chunk of space-time in which they occur. A morphognostic can thus be viewed as a map of progressively larger nested chunks of space-time information forming a hierarchy of contexts that can be used to control responses.

An organism contains a “current” morphognostic which is constantly updated by ongoing sensory inputs from the world, and is thus a working memory representation of the world.

Scaling to prevent information explosion is accomplished by the aggregation feature, which also means that more recent and nearby events are recorded in greater precision than events more distant in space and time.

The following are general definitions of the spatial and temporal morphognostic neighborhoods. The software is parameterized to permit variations of these definitions.

##### Morphognostic spatial neighborhoods

The cellular automaton localizes an elementary neighborhood at a cell:

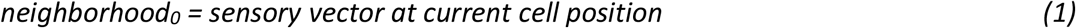

A non-elementary neighborhood consists of an *NxN* set of *sectors* surrounding a lower level neighborhood:

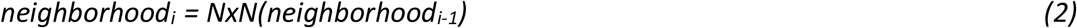

Where *N* is an odd positive number.

The value of a sector is a vector representing an aggregation of the sensory vectors that occur within it:

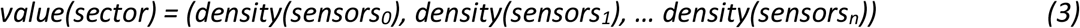

Where *density(sensorsi)* is an aggregation function, e.g. average of the values of the sensory vectors for dimension *i*.

##### Morphognostic temporal neighborhoods

A neighborhood contains events that occur within a *duration*, which is a time window between the present and some time in the past. Here is a possible method for calculating the duration of neighborhood *i*, where the duration grows by a factor of 3:

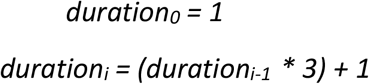

##### Morphognostic example

Figure 2 is an example of a morphognostic structured as a nested set of neighborhoods in a cellular automaton where a cell can have five possible states. On the left side of the figure is a world that contains various cell state values and a mobile organism depicted by the blue triangle. The organism can sense the state of the cell it is positioned on using a 5 dimensional sensory vector, one dimension for each possible cell state value.

**Figure 2 –.**
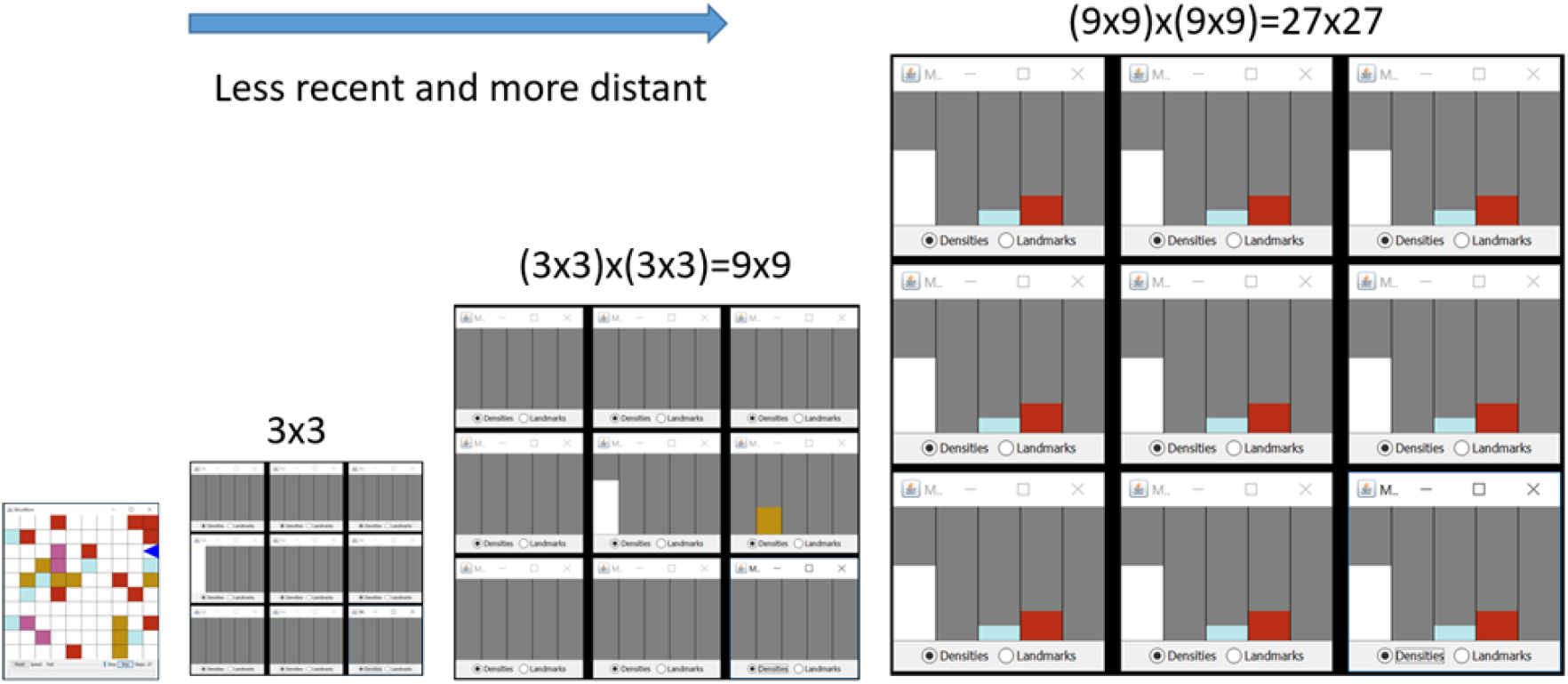
Cellular automaton implementation of Morphognosis.

To the right of the world is a 3×3 morphognostic neighborhood centered on the cell occupied by the organism. The values of each of the 9 sectors in the neighborhood are shown as 5 aggregated sensory vectors input to the organism in the chunk of space-time determined by the sector’s spatial boundaries and the neighborhood’s duration. The aggregation in this example is done by averaging the sensory vector values.

Looking at the left-middle sector of the 3×3 neighborhood, which has a state value of “white” recorded, it can be inferred that the organism sensed the white value of the cell to its left in the previous step, and then moved right to its current position. The 9×9 and 27×27 neighborhoods continue this nesting process to greater spatial and temporal extents.

#### 2.1.2. Metamorphs

In order to navigate and manipulate the environment, it is necessary for an agent to be able to respond to the environment. A *metamorph* embodies a morphognostic→response production rule. A metamorph is acquired from a sensory-response interaction with the world. The sensory input updates the current morphognostic, and when a response is subsequently generated, a metamorph is created to capture the morphognostic→response relationship. The set of metamorphs captures long-term memories of an organism’s interactions with the world.

The set of Metamorphs are used to train the ANN, as shown in Figure 3, to learn the responses associated with morphognostics. A flattening procedure transforms a metamorph’s morphognostic into an ANN input. During operation/testing, the current morphognostic is input to the ANN to produce a response.

**Figure 3 –.**
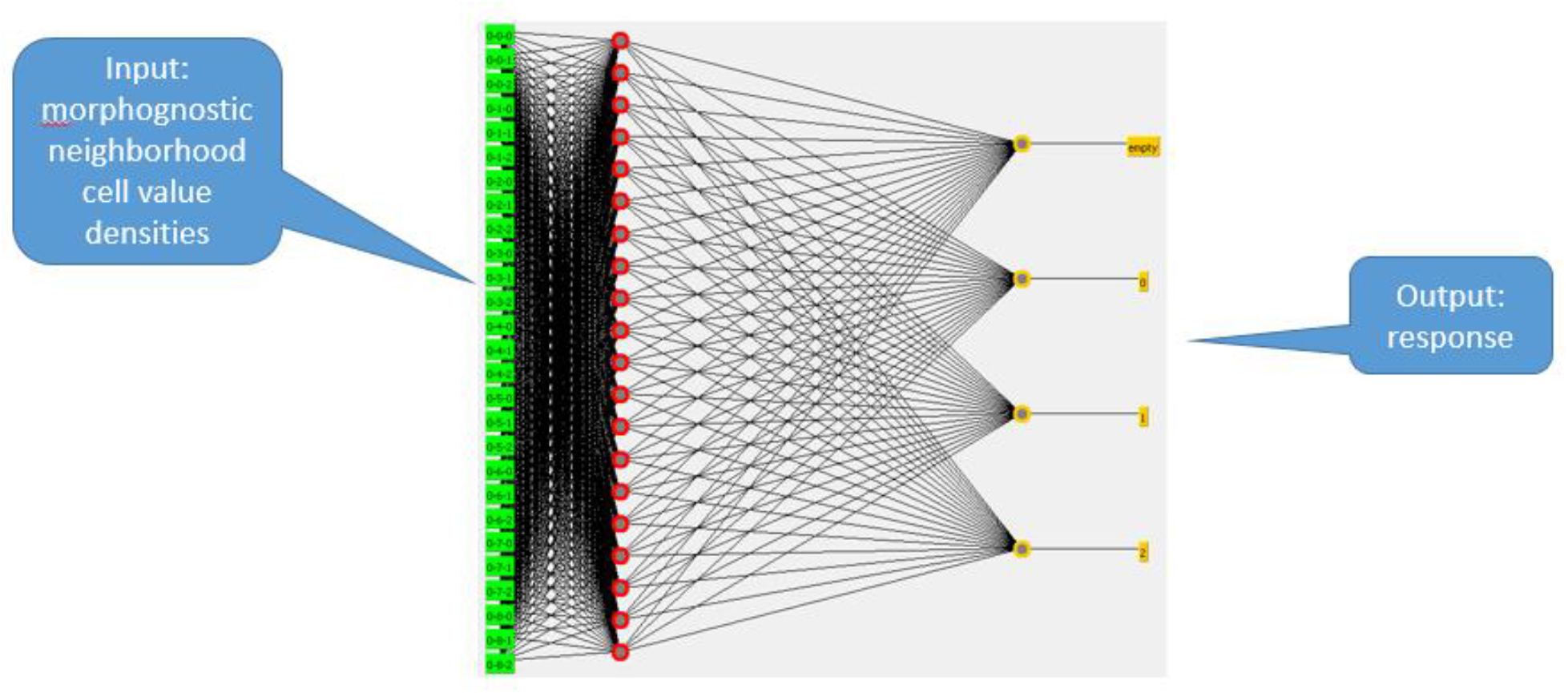
Metamorph artificial neural network.

### 2.2. Honey bees

#### 2.2.1. Behavior

A brief explanatory video is available on YouTube: https://www.youtube.com/watch?v=kUAv2QO7qYM

##### Sensory and response capabilities

The world is stepped by time increments. At each step, a bee is *cycled*. A cycle consists of inputting sensory information and outputting a response that affects the world.

###### Sensory

The world is known by external sensory inputs drawn from its current cell position. A bee also has internal state information that is input to its sensors.

External inputs:

Hive presence.

Nectar presence.

In-hive bee nectar signal: Orientation and distance to nectar.

Internal state:

Orientation.

Carrying nectar.

###### Response

A bee outputs one of the following responses at each step:

Wait,

Move forward,

Turn in compass directions:

N, NE, E, SE, S, SW, W, NW

Extract nectar,

Deposit nectar,

Display nectar distance.

##### World

Figure 4 shows a graphical view that shows a hive (central yellow area), three bees, and three flowers. The topmost flower contains a drop of nectar to which the topmost bee, as best it can in a cellular grid, is indicating the direction and an approximate distance to, as indicated by the orientation of the bee and the length of the dotted line, respectively. The world is bounded by its edges, meaning bees cannot leave one edge and appear on the opposite side. An attempt to move beyond the edge results in a forced random change of direction.

**Figure 4 –.**
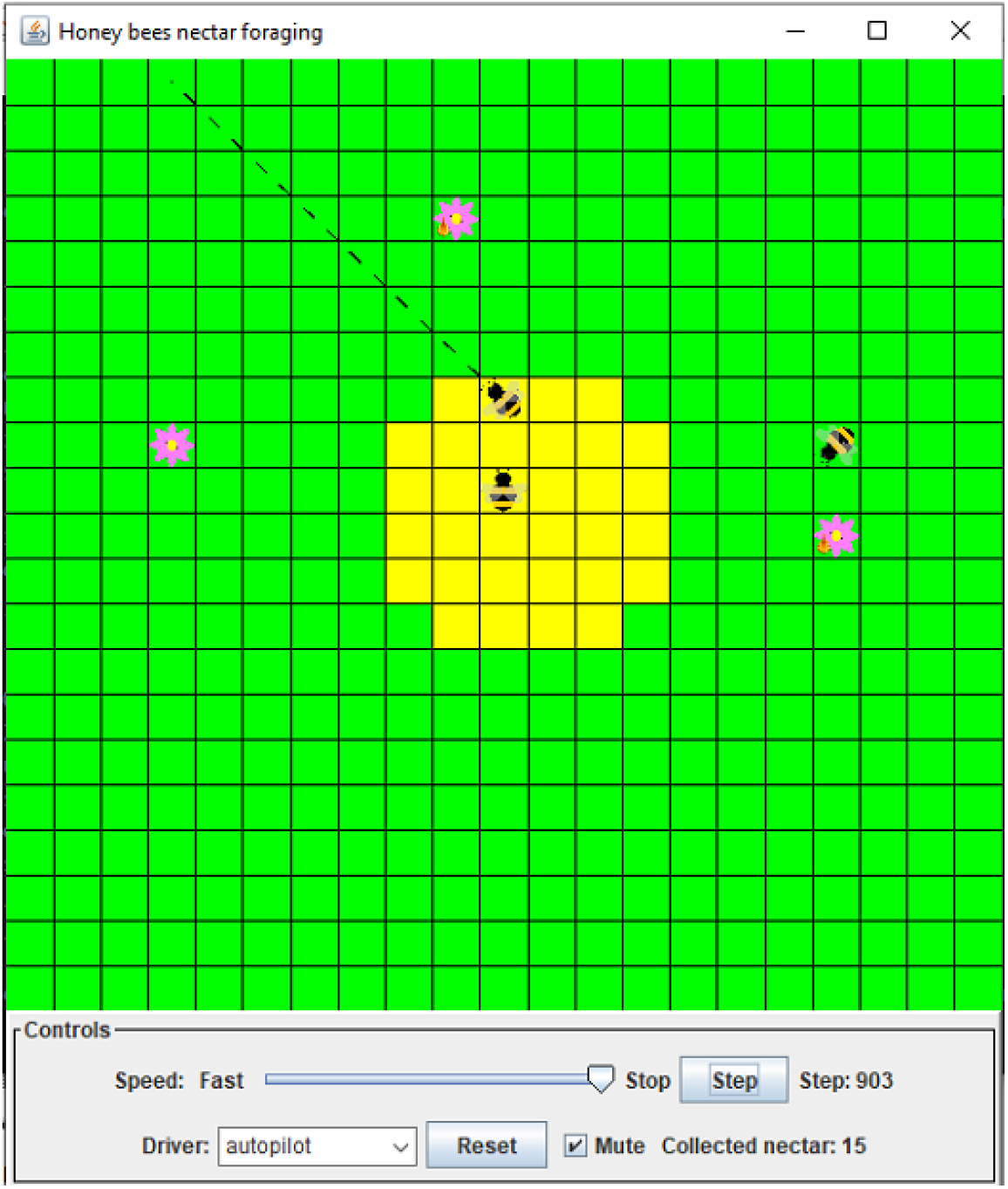
Graphical view.

##### Bees

A bee occupies a single cell and is oriented in one of the eight compass directions and moves in the direction of its orientation. Only one bee is allowed per cell. An attempt to move to an occupied cell is disallowed. If multiple bees move to the same empty cell, a random decision is made to allow one bee to move. Bees can carry a single unit of nectar. Bees are initialized in the hive at random positions and orientations.

##### Flowers

A flower occupies a single cell outside of the hive at a random location. A flower’s cell may also be occupied by a single visiting bee. Flowers are initialized with nectar, which after being extracted by a bee, will probabilistically either replenish after a specific time or immediately replenish. In the latter case, the bee will sense the presence of surplus nectar and will perform a dance to indicate its direction and distance once it returns to the hive. Flowers are initialized at random locations.

##### Foraging

The bees forage in two phases. In phase one, the nectar discovery phase, a bee flies about in a modified Brownian motion until it encounters a flower with nectar. Phase two is a deterministic process that deals with known nectar. Phase two is described below.

Once discovered, the bee extracts the nectar from the flower, flies directly to the hive and deposits the nectar in the hive. If the bee, after depositing the nectar, remembers that the flower contained “surplus” nectar, meaning more nectar than the bee could carry, it will commence a dance which is analogous to the waggle dance in that it indicates the direction and distance to the nectar, but suitable for a grid-world.

The dance is sensed by other bees in the hive, including the dancer. The direction is indicated by orienting toward the nectar. The directions are confined to the eight compass points. The distance is indicated by displaying a value for short or long distance. Both direction and distance can be sensed by bees in the hive. The graphical view draws a short or long dotted line as a visual representation.

Once a bee completes the dance, it and any other bees in the hive that sensed the dance will proceed in the direction of the nectar for the distance exhibited by the dance. If any of these bees encounters nectar en route, it will switch over to extracting the nectar and returning with it to the hive, possibly performing a dance there. If no nectar is encountered en route after traveling the indicated distance, the bee resumes phase one foraging.

If no surplus nectar was sensed after extracting the nectar, the bee will switch to phase one foraging immediately after depositing the nectar.

##### Scenario

Figures 5 through 11 present a graphical nectar foraging scenario.

**Figure 5 -.**
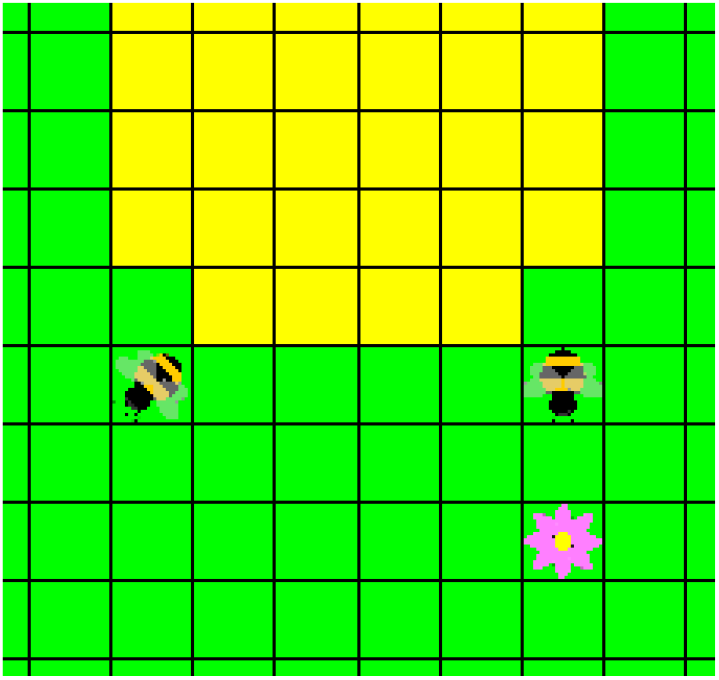
Bee on right is moving down and is about to light on flower.

#### 2.2.2. Implementation

##### Modes

In *autopilot* mode, the bees forage programmatically, meaning the bees are controlled by a program that is hand-written specifically for foraging. The autopilot behavior is optimal, and is thus the goal behavior for learning. Autopilot mode generates metamorphs that are used to train the neural network, as shown in Figure 12. Since phase one foraging consists of semi-random movements, metamorphs are only generated in phase two, dealing with known nectar. Once trained, the bees can be switched to *metamorphNN* mode, in which the neural network drives phase two behavior. Phase one behavior remains semi-random in metamorphNN mode. While in metamorphNN mode, new metamorphs are not accumulated.

**Figure 6 –.**
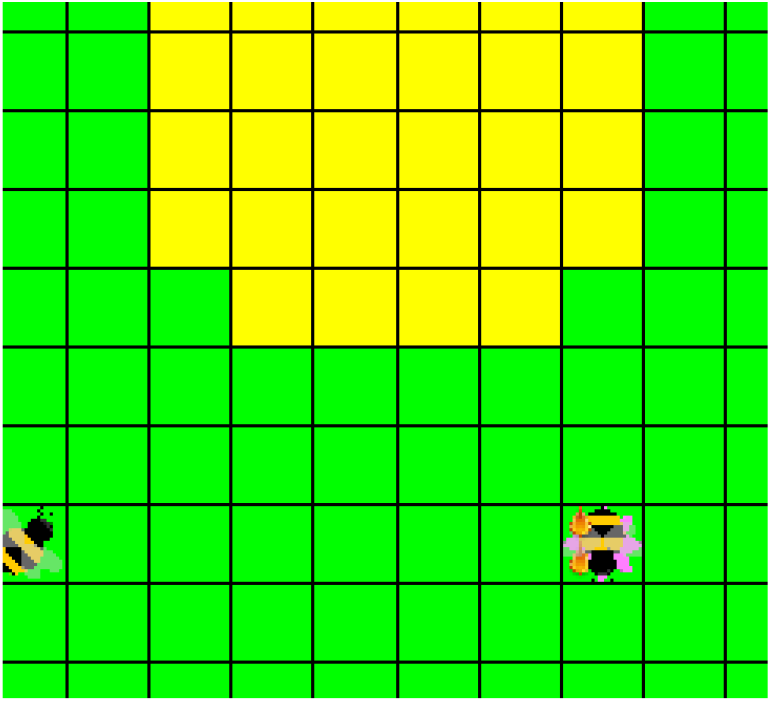
Bee has extracted nectar from flower.

**Figure 7 –.**
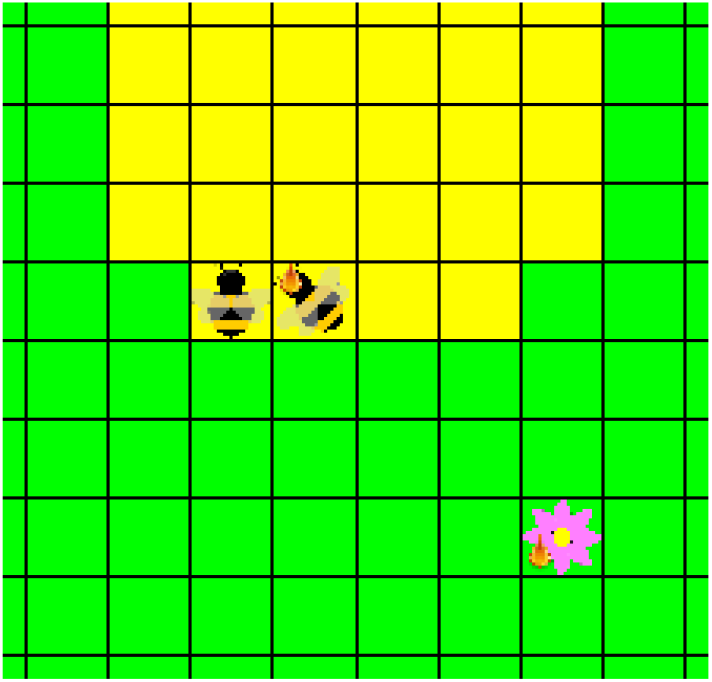
Bee with nectar returns directly to the hive to deposit nectar. It is also aware of surplus nectar remaining in the flower. The other bee is incidentally also in the hive.

**Figure 8 –.**
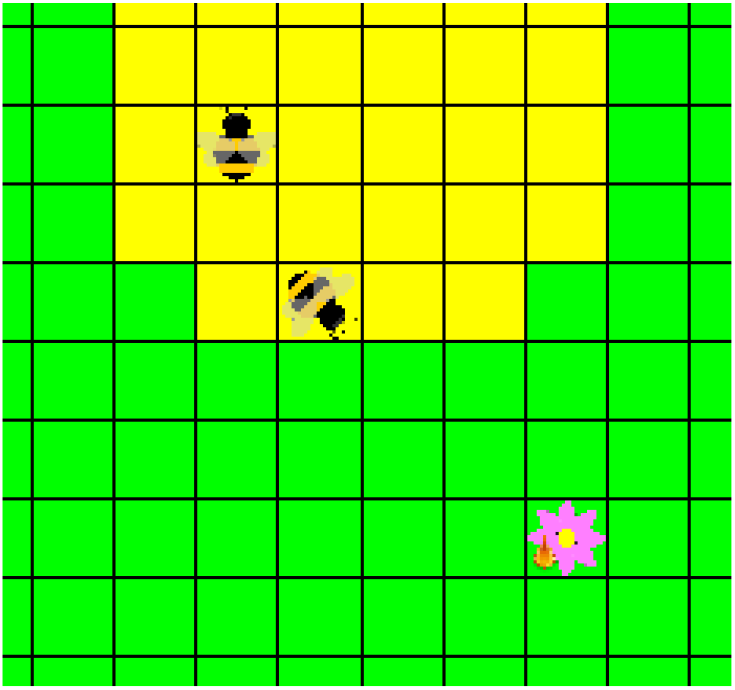
Bee has deposited nectar in the hive. Since the bee knows there is surplus nectar, the bee performs the first part of dance: orient toward nectar. If there was no surplus nectar the bee would resume foraging. The other bee is moving about the hive.

**Figure 9 –.**
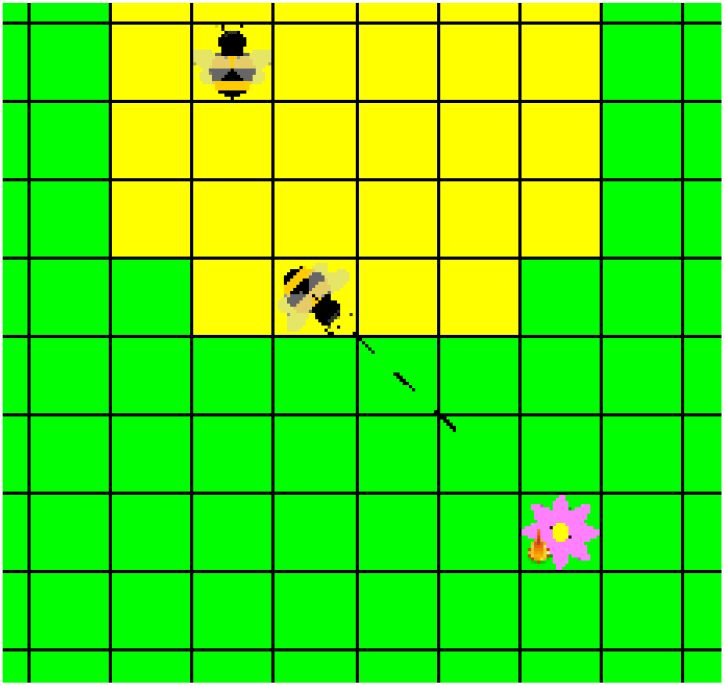
The second part of dance: indicate a short distance to nectar, as shown by the dotted line. The other bee has become aware of the direction and distance to the nectar.

**Figure 10 -.**
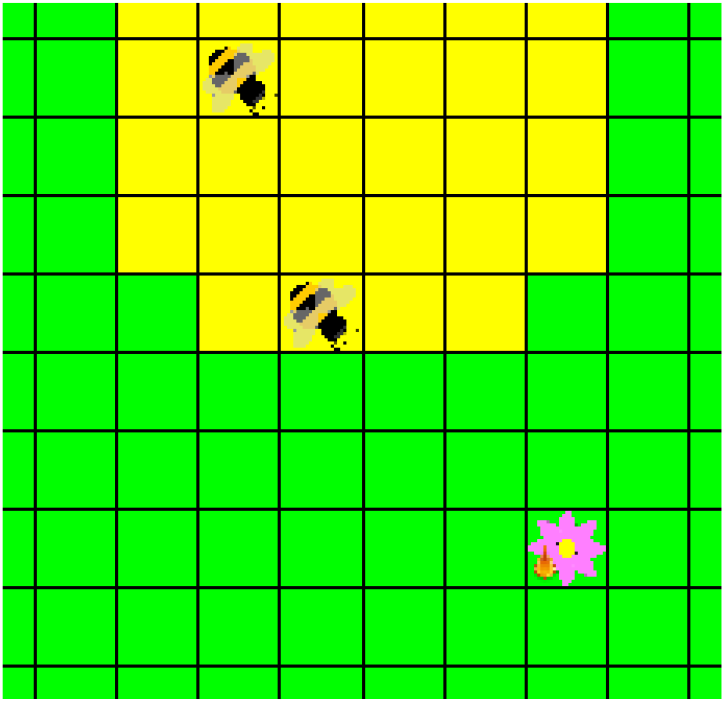
Both bees respond to dance by orienting toward nectar.

**Figure 11 -.**
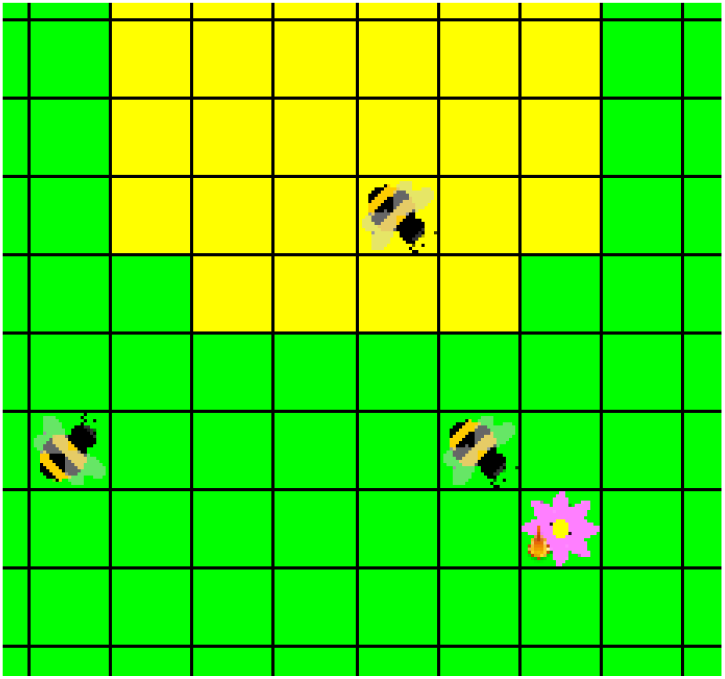
Both bees move toward nectar.

**Figure 12 –.**
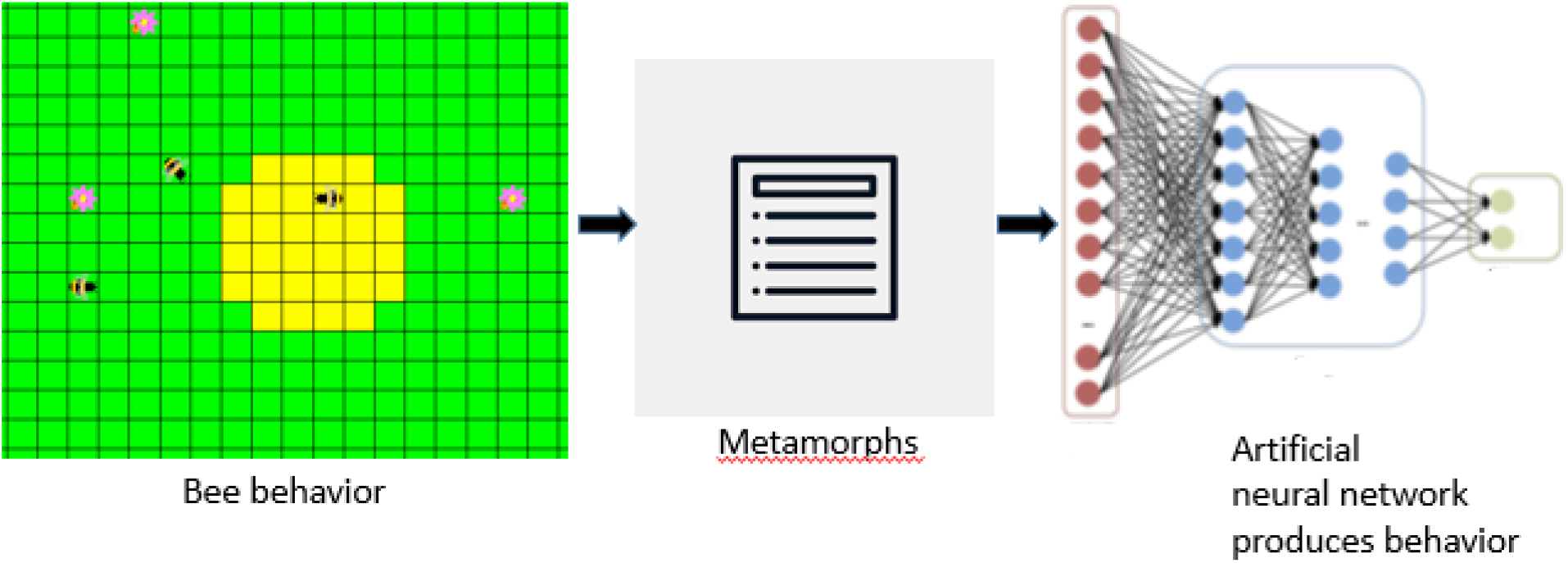
Generating metamorphs to train the neural network.

##### Morphognostic

Each bee contains a morphognostic that maps its sensory inputs as spatial and temporal events that model its state in the environment.

###### Sensory events

There are 22 binary sensory event variables:

0. hive presence
1. nectar presence
2. surplus nectar presence
3. nectar dance direction north
4. nectar dance direction northeast
5. nectar dance direction east
6. nectar dance direction southeast
7. nectar dance direction south
8. nectar dance direction southwest
9. nectar dance direction west
10. nectar dance direction northwest
11. nectar dance distance long
12. nectar dance distance short
13. orientation north
14. orientation northeast
15. orientation east
16. orientation southeast
17. orientation south
18. orientation southwest
19. orientation west
20. orientation northwest
21. nectar carry

###### Neighborhoods

The morphognostic contains 4 3×3 neighborhoods, with durations and sensory event mappings shown in Table 1.

**Table 1.** – Morphognostic neighborhoods.

| Neighborhood | Duration | Sensory events |
| --- | --- | --- |
| 0 | 1 | All except:<br>*surplus nectar presence*<br>*nectar dance distance long*<br>*nectar dance distance short* |
| 1 | 7 | *hive presence*<br>*surplus nectar presence*<br>*nectar dance short distance* |
| 2 | 10 | *hive presence*<br>*surplus nectar presence*<br>*nectar dance long distance* |
| 3 | 75 | *hive presence* |

Neighborhood 0 maps “immediate” events, such as orientation, that are of use only in the present, as denoted by the duration of 1.

Neighborhood 1 has a duration, 7, that allows a bee to retain knowledge of the presence of surplus nectar and/or observation of a dance indicating a short distance. The *nectar dance short distance event*, for example, allows the bee to “count” steps towards surplus nectar. When the event expires due to the duration of the neighborhood it no longer affects the bee’s behavior.

Neighborhood 2 serves the same purpose as neighborhood 1, except for *nectar dance long distance* event, for which the duration, and thus steps, is greater than for the *nectar dance short distance* event.

Neighborhood 3, as well as all the other neighborhoods, track the presence of the hive as it is recorded in its 3×3 sectors for a long duration of 75. This allows the bee to locate the hive after possibly lengthy foraging and return with nectar. On the rare occasion that 75 steps are taken without returning to the hive, its location will be lost and the bee will be forced to return to the hive without nectar.

Morphognostic neighborhoods can be configured to either keep an average density value over its duration, or an on/off value, meaning the sensory event value is 1 if the event occurs at any time within the neighborhood’s duration window. Although surrendering information, the on/off configuration is chosen for the honey bees to improve training time while retaining acceptable performance.

###### Example

Figures 13a and 13b show the state of the bee selected by the red square for neighborhood 2 of its morphognostic.

**Figure 13a –.**
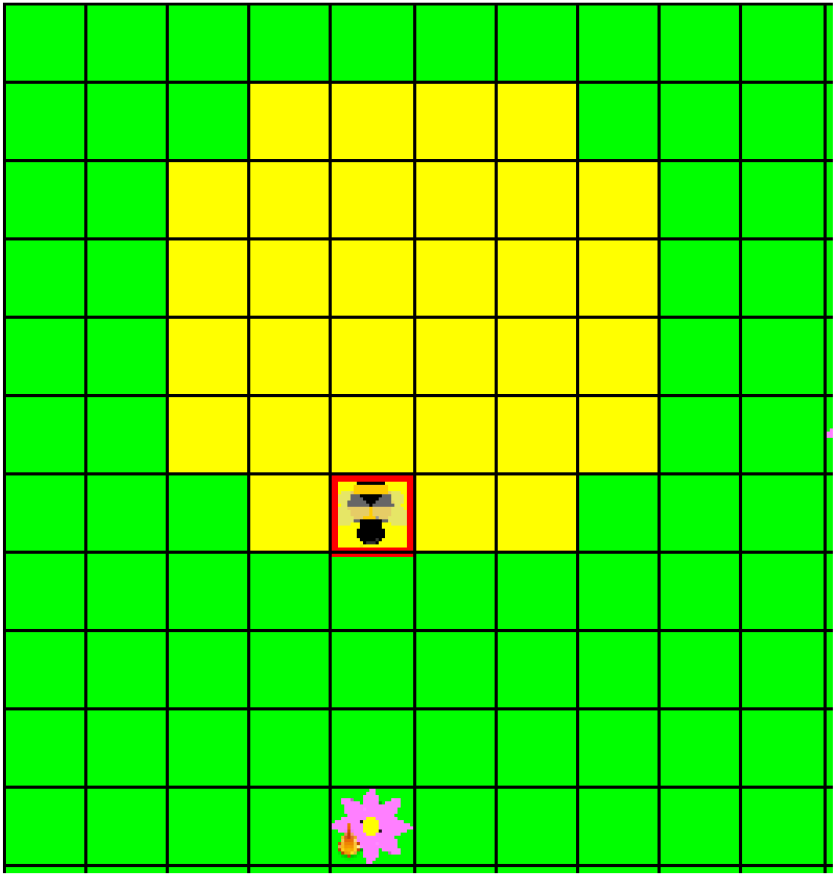
Bee after dance indicating surplus nectar. The next step is to proceed toward nectar.

**Figure 13b –.**
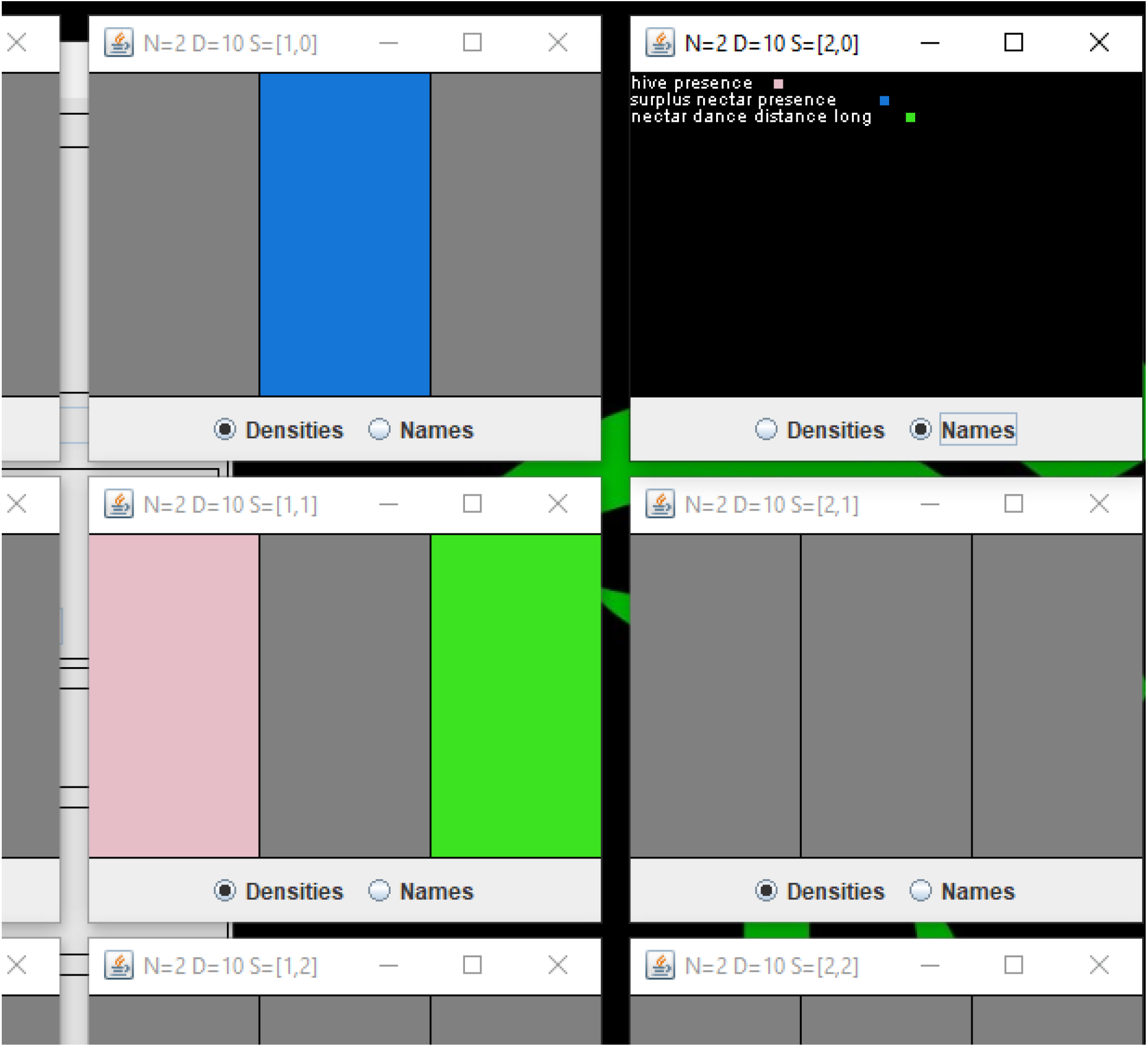
Morphognostic neighborhood 2. At the center sector [1, 1] the *hive presence* and *nectar dance distance long* sensory events are recorded. The location of the surplus nectar is recorded in sector [1, 0] and was used to orient toward the surplus nectar as part of the dance.

##### Code

The Java code is available on GitHub: https://github.com/morphognosis/HoneyBees

## 3. Results

### 3.1. Artificial neural network

The artificial neural network used was the MultiLayerPerceptron class in the Weka 3.8.3 machine learning library (https://www.cs.waikato.ac.nz/ml/weka/).

These parameters were used:

- learning rate = 0.1
- momentum = 0.2
- training epochs = 5000

The morphognostic configured as previously described, four 3×3 neighborhoods, produces 234 binary inputs to the network. There are 14 outputs representing the honey bee responses.

### 3.2. Base level testing

Neither a randomly generated responses nor an untrained network resulted in any nectar collected over 20,000 steps in a 3 flower and 3 bee configuration.

### 3.3. Test flower and bee quantities

In order to determine how the system scales up, three variations of flowers and bees were tested: 3 flowers and bees, 5 flowers and bees, and 7 flowers and bees. The amount of nectar collected was used as a success metric.

The world was set at 21×21 cells, and the hive at radius 3. Flowers were initialized with nectar at random locations outside of the hive. Bees were initialized randomly in the hive. The network was configured with 50 hidden neurons. Running the world for 20,000 steps on autopilot generated a metamorph dataset to train the neural network on. Datasets were generated for 10 trials.

Table 2 shows the average training dataset size and training accuracy. Of note is the increase in the number of metamorphs as the world become more complex with additional flowers.

**Table 2.** – Number of metamorphs and training accuracy by varying flower and bee quantities.

| Flowers and bees | Metamorphs | Accuracy |
| --- | --- | --- |
| 3 | 972.4 | 99% |
| 5 | 2271.7 | 99% |
| 7 | 3852.3 | 99% |

Figure 14 shows the results of running optimally (Autopilot) vs. with the trained network (Morphognosis). The network performs comparably.

**Figure 14 –.**
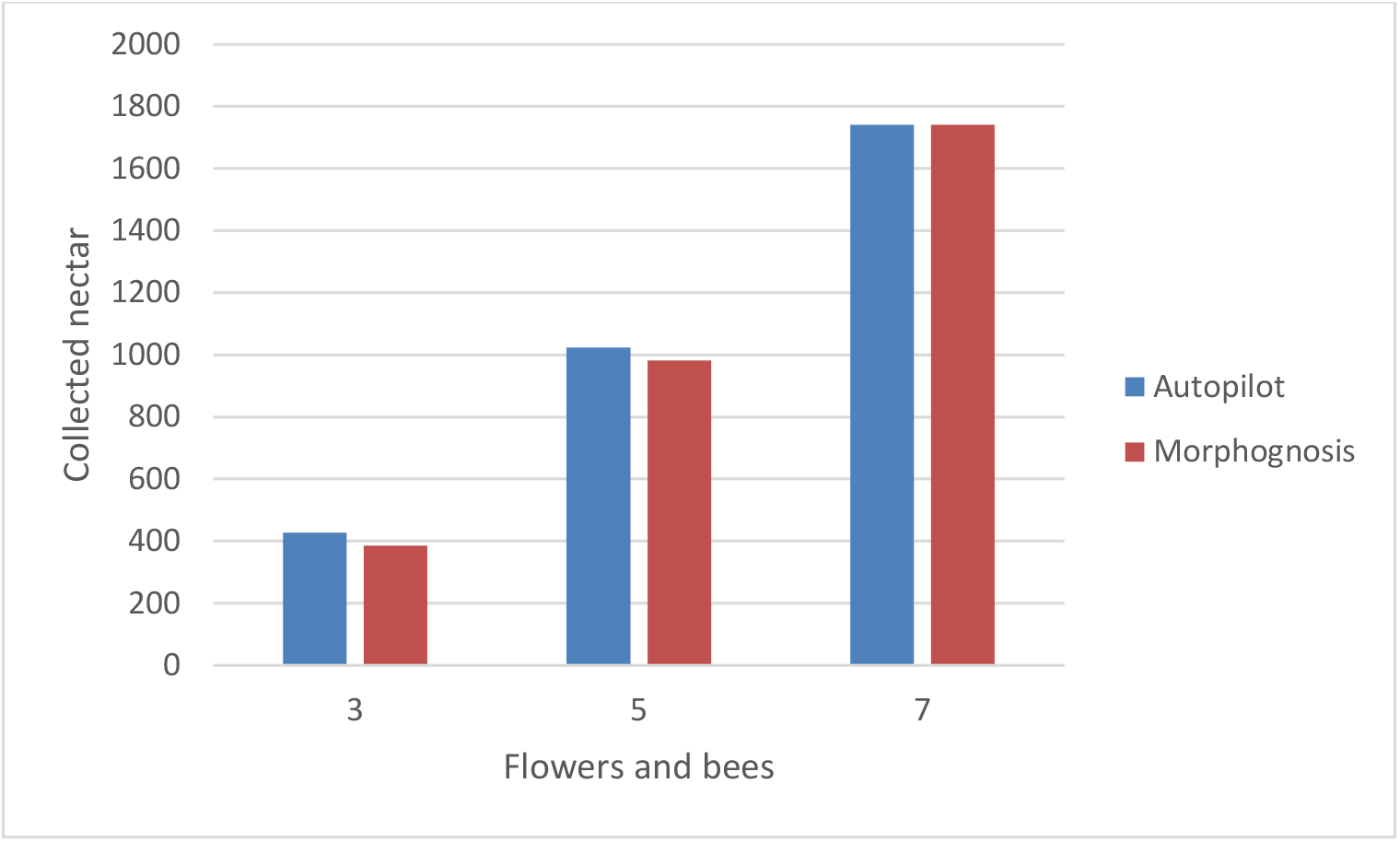
Collected nectar for variations of flowers/bees.

### 3.4. Test hidden neurons

In order to observe how the system is affected by the neural network size, three variation of hidden neuron quantities were tested: 25, 50, and 100.

Table 3 shows the average training dataset size and training accuracy.

**Table 3.** – Number of metamorphs and training accuracy by varying hidden neurons.

| Hidden neurons | Metamorphs | Accuracy |
| --- | --- | --- |
| 25 | 913.9 | 99% |
| 50 | 972.4 | 99% |
| 100 | 887.4 | 99% |

Figure 15 shows the results, indicating that fewer hidden neurons are sufficient to achieve comparable performance.

**Figure 15 –.**
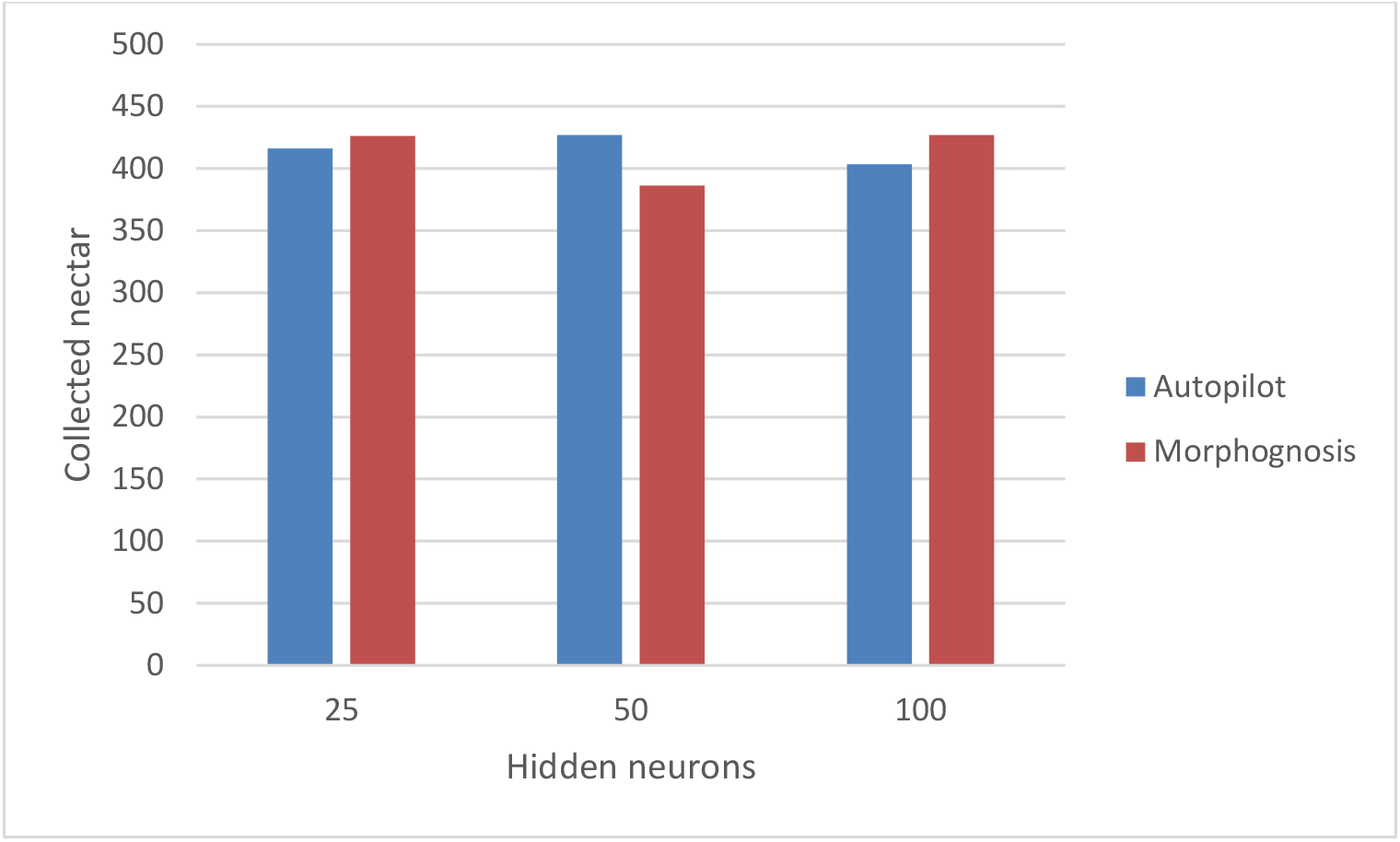
Collected nectar for variations of hidden neurons.

### 3.5. Test hive radius

In order to observe how the system is affected by the hive size, two variation of hive sizes were tested: radii of 2 and 3.

Table 4 shows the average training dataset size and training accuracy. Of note is the reduction in metamorphs with a smaller hive. This is likely due to fewer “trajectories” to and from the hive.

**Table 4.** – Number of metamorphs and training accuracy by varying hive radius.

| Hive radius | Metamorphs | Accuracy |
| --- | --- | --- |
| 2 | 622.4 | 99% |
| 3 | 972.4 | 99% |

Figure 16 shows the results, indicating that a smaller hive reduces the amount of nectar collected. A possible contributing factor for this is congestion due to bee collisions.

**Figure 16 –.**
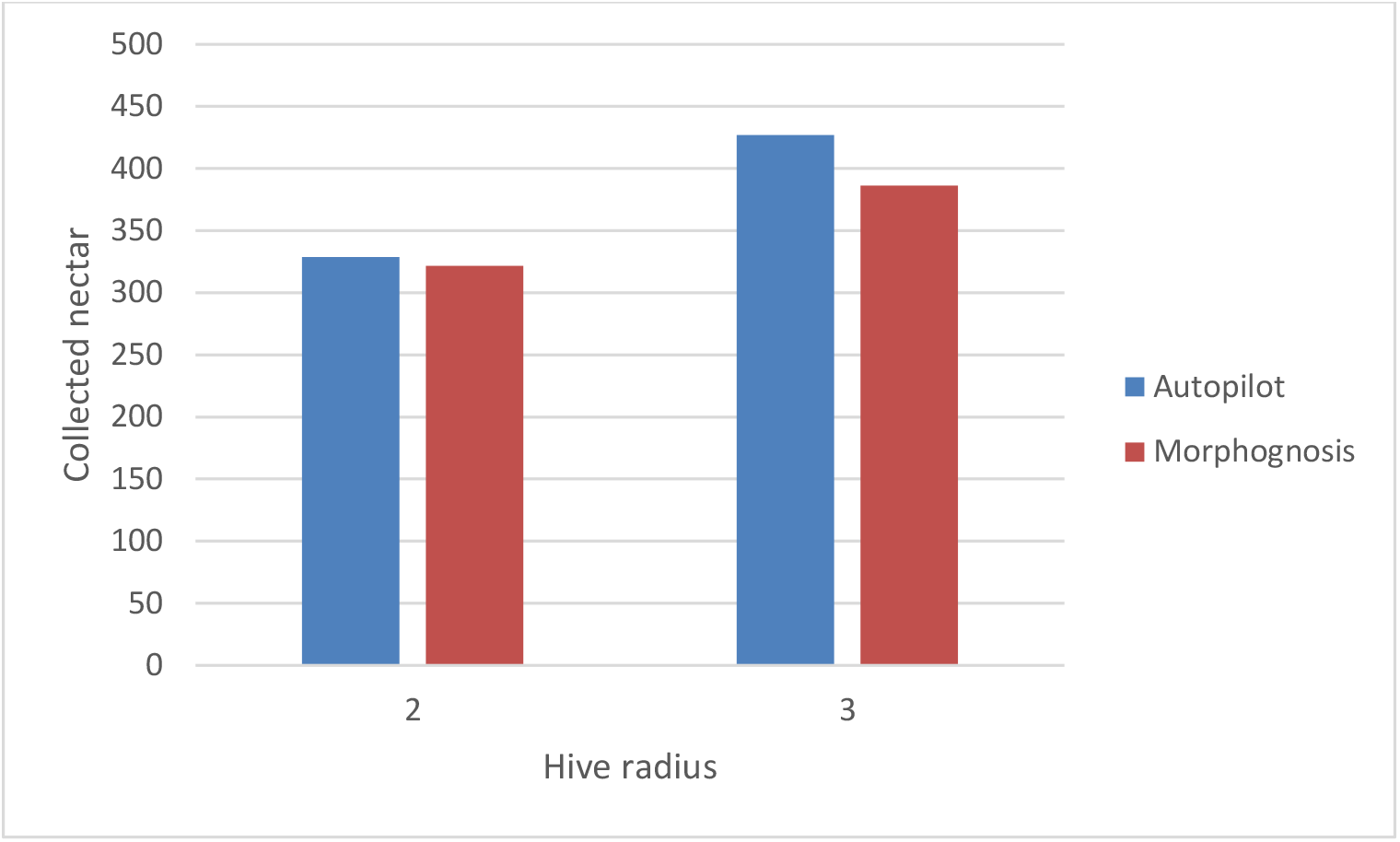
Collected nectar for variations of hive radius.

### 3.6. Recurrent neural network performance

A key ability of a honey bee is to be able to track the location of the hive as it forages. This allows it to return to the hive with nectar. While it is known that a recurrent neural network (RNN) is capable of learning positions along a specific path (Cueva and Wei 2018), the honey bee foraging task requires the ability to perform dead-reckoning (also known as path integration) over many possible paths.

To test the ability of an RNN to learn a general dead-reckoning task, a Long Short Term Memory (LSTM) recurrent network (Hochreiter and Schmidhuber 1997) was trained given sequences between 5 and 15 steps consisting of random orientation changes and forward movements probabilistically identical to those used by the honey bees. The output is the direction to the starting position. Despite variations in the network capacity, the training accuracy averaged approximately 30%, which was about the same as a random guess.

An LSTM was also tested on the deterministic portion of the foraging task. This is the activity that occurs after a nectar source is discovered. This consists of extracting the nectar, flying to the hive location, and depositing the nectar. If surplus nectar was sensed before leaving the flower, a dance must also be performed indicating the distance and direction to the nectar, following which the bee sets out in the direction of the nectar.

To implement this, the network is provided a sensory map marking the location of the hive when nectar is discovered, which is sufficient to allow the bee to fly back to the hive, and to indicate the direction to the surplus nectar if a dance is necessary. This marking must be learned by the network, as it only occurs at the beginning of the forage. Training with an equal mixture of dance and non-dance forages, 100% of forages were successfully learned. However, when the RNN was tested with flowers spatially displaced from their training locations, performance decreased dramatically. For example, a displacement of only three cells resulted in 0% success. In contrast, the overall performance of the Morphognosis network was unaffected by this displacement. This is due to the hive location being recorded at multiple levels in its memory hierarchy at varying degrees of granularity. This flexibility is important for a simulation of honey bees that forage in a world of variable flower locations.

It is also important to note a distinction between the training and testing regimens of RNNs and Morphognosis. RNNs are trained with batches of sequences. Each sequence, possibly having a variable length, has a beginning and end. A test inputs a sequence to the trained network for classification and prediction. Morphognosis, in contrast, having its temporal (and spatial) state embedded in the input, is not bounded by sequences: a honey bee generates a set of training metamorphs as it forages continuously, with no delimiting breaks. This more closely resembles an animal learning situation in nature.

The LSTM network used is in the Keras 2.2.4 machine learning package: https://keras.io/

## 4. Conclusion

The brain, a complex structure resulting from millions of years of evolution, can be viewed as a solution to problems posed by an environment existing in space and time. Internal spatial and temporal representations allow an organism to navigate and manipulate the environment. Following nature’s lead, Morphognosis comprises an artificial neural network enhanced with a framework for organizing sensory events into hierarchical spatial and temporal contexts.

It has been demonstrated that using the augmenting facilities provided by Morphognosis, an artificial neural network is capable of performing the honey bee foraging and dancing task. It has also been demonstrated that without this augmentation, an artificial neural network performs poorly at both path integration and returning nectar to the hive when flower locations are displaced.

The successful simulation of honey bee foraging behavior suggests future projects are worth undertaking:

- The metamorph structure bears a close resemblance to deep reinforcement learning training elements (Francois-Lavet et al. 2018), suggesting the possibility of applying such learning to implement goal-seeking behavior.
- The aggregation scheme that supports scalability is a simple histogram-like method for dimensionality reduction.
  - The use of ANN dimensionality reduction techniques, such as autoencoding, might scale with higher information content.
  - The value of each neighborhood sector essentially represents a single centroid of sensory event values that have occurred in its space-time cube. An extension of this would be to retain multiple centroids within a sector, possibly weighted by frequency, increasing in number for higher level neighborhoods which encompass greater extents of space-time. This might increase the richness of behavioral variability while limiting information overload.
- The model is currently implemented in a cellular automaton spatial grid of cells. However, it is not inherently tethered to this platform and in fact may benefit from extending beyond it.
- The configuration of the morphognostic is vital to successful performance. For the honey bee task, this was a manual design. This process should be amenable to optimization/evolution methods.

